# Characterizing neuronal synaptic transmission using stochastic hybrid systems

**DOI:** 10.1101/582445

**Authors:** Zahra vahdat, Zikai Xu, Abhyudai Singh

## Abstract

Action potential-triggered release of neurotransmitters at chemical synapses forms the key basis of communication between two neurons. To quantify the stochastic dynamics of the number of neurotransmitters released, we investigate a model where neurotransmitter-filled vesicles attach to a finite number of docking sites in the axon terminal, and are subsequently released when the action potential arrives. We formulate the model as a Stochastic Hybrid System (SHS) that combines three key noise mechanisms: random arrival of action potentials, stochastic refilling of docking sites, and probabilistic release of docked vesicles. This SHS representation is used to derive exact analytical formulas for the mean and noise (as quantified by Fano factor) in the number of vesicles released per action potential. Interestingly, results show that in relevant parameter regimes, noise in the number of vesicles released is sub-Poissonian at low frequencies, super-Poissonian at intermediate frequencies, and approaches a Poisson limit at high frequencies. In contrast, noise in the number of neurotransmitters in the synaptic cleft is always super-Poissonian, but is lowest at intermediate frequencies. We further investigate changes in these noise properties for non-Poissonian arrival of action potentials, and when the probability of release is frequency dependent. In summary, these results provide the first glimpse into synaptic parameters not only determining the mean synaptic strength, but also shaping its stochastic dynamics that is critical for information transfer between neurons.

## 1 INTRODUCTION

We present a mechanistic stochastic model to characterize the statistics of the number of neurotransmitters released at a neuronal synapse. While there is rich literature regarding synaptic transmission as a deterministic process [1–5], increasing evidence points to diverse noise mechanisms at play during this process [6–10]. Counterintuitively, noise has been shown to sometimes enhance signaling and information transfer between neurons [11–16].

Stochastic Hybrid Systems (SHS) constitute an important class of mathematical models that integrate discrete stochastic events with continuous dynamics. Given their generality and scope, SHS have been successfully used for modeling stochastic phenomena in a variety of biological processes [17–33], including neuronal dynamics [34, 35]. Building up on our previous work [36], we consider the SHS framework for modeling the neurotransmitter dynamics. More specifically, the model consists of *M* ∈ {1, 2,…} docking site at the axon terminal (Fig. 1). Neurotransmitter-filled vesicles attach to these docking sites with a given probabilistic rate that is proportional to *M* −***n***, where ***n*** is the number of already docked vesicles. Action Potentials (APs) arrive at a given frequency, such that, the time between two successive APs follows an arbitrary positively-valued probability density function *g*. Each docked vesicle has a certain probability of release, and based on this probability, APs cause a fraction of the docked vesicles to release their neurotransmitter content into the synaptic cleft (gap between neurons). Finally, released neurotransmitters decay (or are removed) at a constant rate. Details of the model along with sample stochastic realizations are shown in Fig.1.

**Figure 1:**
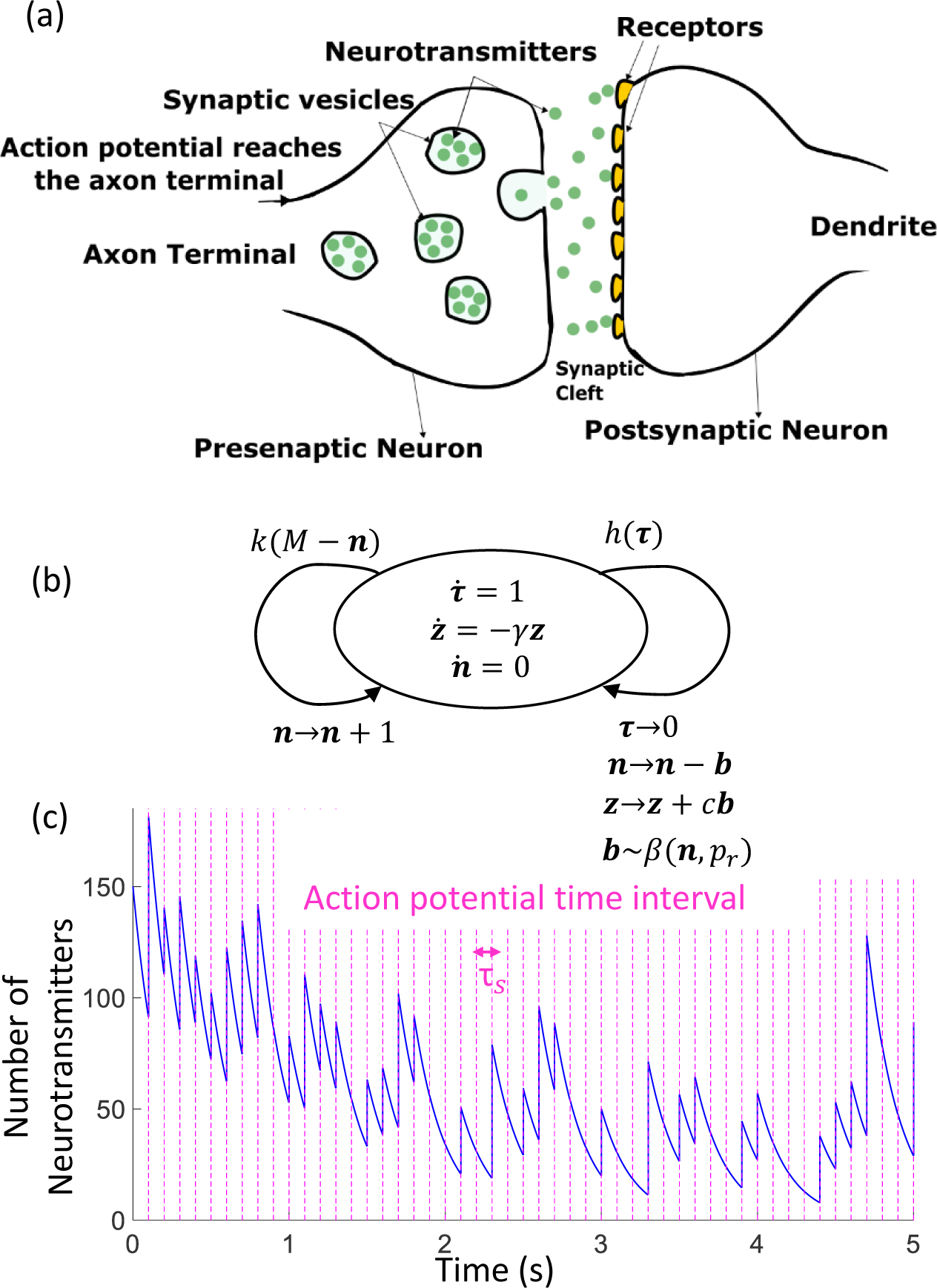
**(a): Illustration of a chemical synapse.** Neurons communicate via synaptic transmission, where APs in the presynaptic neuron trigger release of neurotransmitters into the synaptic cleft. **(b): Stochastic Hybrid Systems model of neuronal synaptic transmission.** Two families of random resets characterize the SHS: the first is the arrival of APs based on an internal timer that measures the time elapsed from the previous AP. As shown in (3), this can be used to model any probability distribution for the inter-arrival time. The second reset is the creation of new release-ready vesicle that happens with rate *k*(*M −****n***), where *M* is the maximum possible number of docking sites, *k* is the refilling rate per site. If ***n*** is the number of docked vesicles, then *M −****n*** is the number of empty sites. **(c): Sample realization for the number of neurotransmitters in the synaptic cleft.** For this plot we assumed deterministic arrival of APs with inter-arrival time ***τ***_***s***_ = 0.1 *sec, k* = 1 *sec*^−1^, *γ* = 5 *sec*^−1^ and probability of release *p*_*r*_ = 0.5. The number of neurotransmitters in each vesicle is assumed to be *c* = 30.

The level of neurotransmitters in the synaptic cleft ultimately determines the downstream biological activity through binding and opening of ion channels in the postsynaptic neuron. How does noise in the neurotransmitter level depend on the frequency of AP arrivals, the probability of vesicular release, and other synaptic parameters? To address this question, we derive exact analytical results for the statistical moments of the SHS state space. While solving moment dynamics for SHS often suffers from the problem of moment closure [37–39], here we find exact analytical formulas for the steady-state mean and noise levels for both the number of neurotransmitters released per AP, and the neurotransmitter level in the synaptic cleft.

Analysis of these formulas reveals that depending on the parameter regime, the noise in the number of vesicles released can vary monotonically or non-monotonically with the arrival frequency. In the latter case, noise is sub-Poissonian at low frequencies, super-Poissonian at intermediate frequencies, and approaches a Poisson limit at high frequencies. In contrast, noise in the number of neurotransmitters in the synaptic cleft is always super-Poissonian, but is lowest at intermediate frequencies. The paper is organized as follows: in Section II we recast synaptic transmission models in the SHS framework, and systematically analyze this class of stochastic models in Section III to derive exact moment dynamics. Based on these results, we present our key findings in Section IV, followed by the Conclusion section.

## 2 Model Formulation

Fig.1(a) shows the structure of a (chemical) synapse that permits communication between neurons via released neurotransmitters. Briefly, this process is triggered when an electrical stimulus or Action Potential (AP) reaches the axon terminal of the presynaptic neuron. Inside the axon terminal, vesicles filled with neurotransmitters are synthesized, and they become release-ready by docking close to the cell membrane. Arrival of an AP leads to opening of calcium ion channels, and the influx of calcium causes these docked vesicles to release their neurotransmitter content in the synaptic cleft (extracellular space between the two neurons). The vesicular release process is known to be probabilistic, with each docked vesicle having a certain probability of release. Released neurotransmitters bind to receptors on the cell membrane, triggering an AP in the postsynaptic neuron.

We model the stochastic dynamics of synaptic transmission by a SHS shown in Fig.1(b). The SHS state-space ***x*** = [***n z***]^*T*^, where ***n***(*t*) is the number of docked vesicles, and ***z***(*t*) is the number of neurotransmitters in the cleft at time *t*. Two families of resets occur randomly over time that impact the time evolution of ***x***. The first reset is the arrival of an AP that results in the release of vesicles, and the second reset is the replenishment of the vesicle pool from newly synthesized vesicles. We describe these resets in further detail below, starting with the AP arrival process.

### 2.1 Timer-based arrival of Action Potentials

We assume that APs arrive at times ***t***_*s*_, *s* ∈ {1, 2, 3,…}, such that the time intervals ***τ***_*s*_*≡* ***t***_*s*_ *−****t***_*s−*1_ are independent and identically distributed random variables following an arbitrary positively-valued continuous probability density function (pdf) *g*. To model the timing of APs, we introduce a timer ***τ*** that linearly increases over time

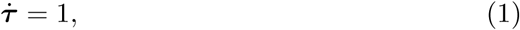

and resets to zero

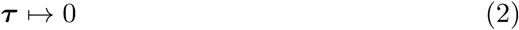

whenever an AP arrives. The arrival of the next AP depends on the state of the timer, and this introduces memory in the arrival process. More specifically, the probability that the AP arrives in the next infinitesimal time interval (*t, t* + *dt*] is given by *h*(***τ***)*dt*, where the *hazard rate*

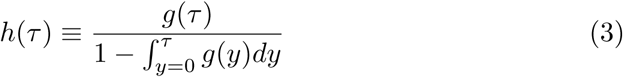

[40–42]. Defining the arrival of events as per (3) ensures that ***τ***_***s***_ follows the pdf *g*

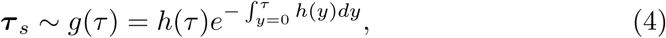

and the corresponding pdf of ***τ*** is given by

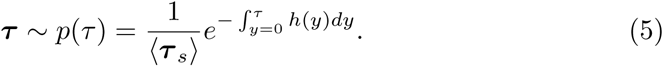

For example, if *g* is exponentially-distributed with mean ⟨***τ***_***s***_,⟩ then *h*(**τ**) = 1/⟨*τ*_*s*_⟩ would be a constant corresponding to a Poisson arrival of events.

Having modeled the timing of the AP arrival process, we next describe its impact on the SHS state space. We assume that each vesicle has a certain probability of release (*p*_*r*_). Thus, an AP causes the SHS state space to jump as per the following reset

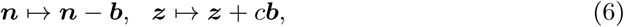

where ***b*** is the number of vesicles released and follows a Binomial distribution

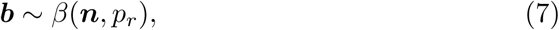

and *c* is the number of neurotransmitters in each vesicle. The reset (6) can be written in terms of ***x*** as

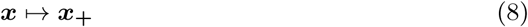

where random variable ***x*_+_** is the state of system immediately after an AP. Using (7) and (8), the mean and covariance matrix of ***x*_+_** is given by

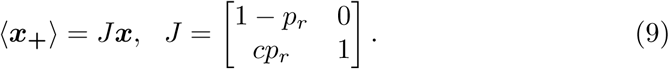

and

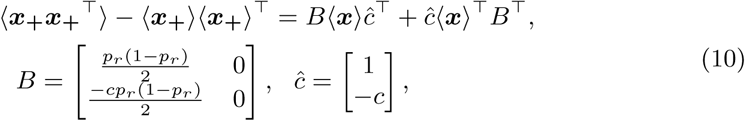

respectively. Note that the stochastic jumps in ***x*_+_** are state-dependent due to the binomial nature of the release process [43, 44].

### 2.2 Stochastic hybrid system model

In addition to timer-based arrival of APs, we also have a second family of resets corresponding to replenishment of vesicles. We assume that the docked-vesicle pool replenishes with a rate *k*(*M −****n***), where *M* is the total number of docking sites (i.e., maximum possible number of vesicles), and *k* represents the refilling rate per empty site. We rewrite this rate in terms of ***x*** as

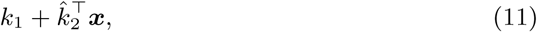

where 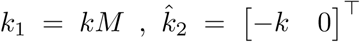. The replenishment process is assumed to occur independently of APs. In the stochastic sense, the probability that a replenishment event occurs in the next time interval (*t, t* + *dt*] is given by *k*(*M −* ***n***)*dt*, and whenever this event occurs

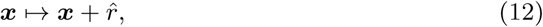

where 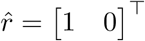 and corresponds to ***n*** increasing by one. In between the two families of resets, the state evolves continuously via a linear system

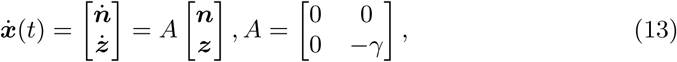

where *γ* is the constant rate of removal or decay of released neurotransmitters.

In summary, we have defined a SHS model for synaptic transmission with continuous dynamics (13), two family of rests that occur with rates (3) and (11), and when they occur, the state space resets via (8) and (12), respectively. It is important to point out that this model with just the timer-dependent reset has been referred to in literature as time-triggered SHS [45–48]. Allowing for a second family of resets that occurs at a rate linearly dependent on ***x*** is a novelty of this work.

## 3 Solving moment dynamics

Next, we derive exact analytical formulas for the first- and second-order momen<u>ts</u> of ***x*** = [***n z***]^*T*^. In this sections we use ⟨ ⟩ to denote the expected value, and 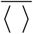 to show the expected value at steady-state.

### 3.1 First-order moment

As shown in Appendix, in between two successive AP arrivals, the conditional moment ⟨***x****|**τ***⟩ evolves as

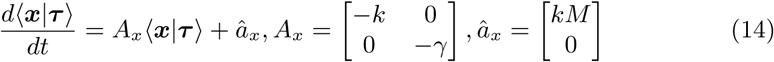

[49]. By solving (14) we have

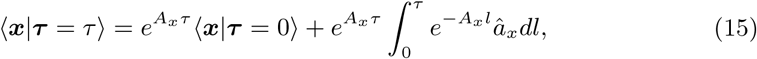

where ⟨***x****|**τ*** = 0⟩ is the expected value just after AP arrival. Given that the time interval between two APs is ***τ***_*s*_ and the reset in mean levels (9), then at steady-state

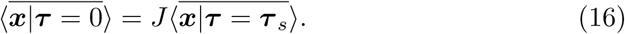

Using (15) for ***τ***= ***τ***_*s*_ together with (16) solves 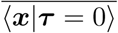, and then using (5) to make ⟨***x****|**τ***⟩ unconditioned with respect to ***τ***, yields

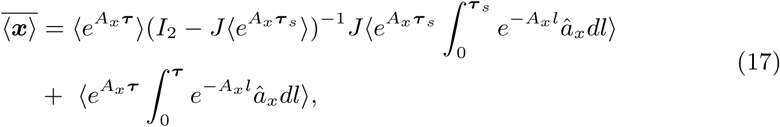

where *I*_2_ is an identity matrix of size 2. Additionally, the system is stable if eighenvalues of 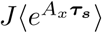 are inside the unit circle. Using all the constant vector and matrices defined in (9)-(12), we can solve for the mean number of docked vesicles

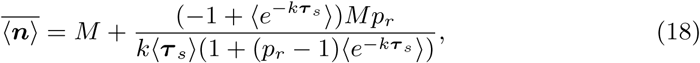

and the mean neurotransmitters level

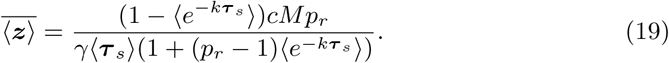

### 3.2 Second order moment

To obtain the second-order moments we define a new vector

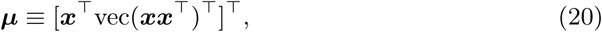

where vec(***xx***^*T*^)∈*ℝ*^*n×*1^ is a vector representation of the matrix ***xx***^*T*^∈*ℝ*^*n×n*^. The new vector ***µ*** =[***n z n***^***^2^***^***nz z n z***^***^2^***^]^T^ contains all the first and second order moments. It can be shown after some algebraic steps (see also Appendix),

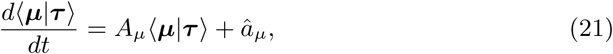

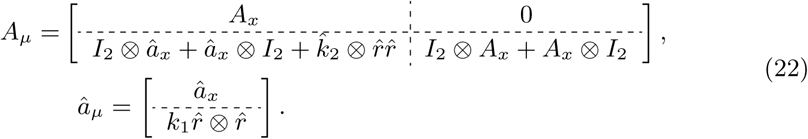

where *⊗* is Kronecker product. Solving (21) gives

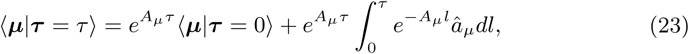

Similar to (16), based on the resets (9) and (10) we can write

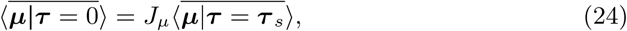

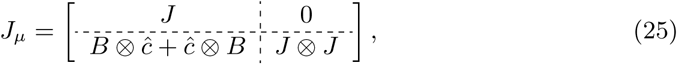

and then using an approach analogous to the first order moment results in

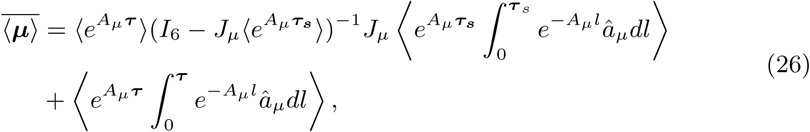

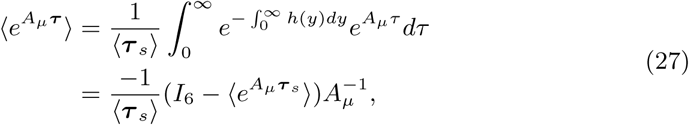

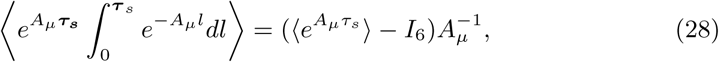

where *I*_*6*_ is an identity matrix of size 6. The eighenvalues of 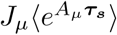 should be inside the unit circle in order for the system to be stable.Using (26) and the first order moments (18) and (19), we obtain the second order moments (43) and (44). Due to the size of these formulas they are presented at the end on the last page.

Throughout the paper we use the steady-state Fano factor (variance over mean) defined by

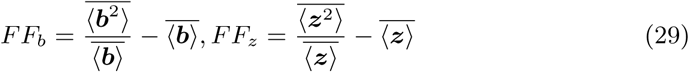

as a metric to quantify noise. Here, *FF*_*b*_ and *FF*_*z*_ denote the Fano factor for the number of vesicles released, and the Fano factor for the number of neurotransmitters, respectively. While *FF*_*z*_ can be calculated using the first and second order moments in (19) and (44), to obtain *FF*_*b*_ we recall that given ***n***, ***b*** *∼ β*(***n***, *p*_*r*_) follows a binomial distribution. Hence,

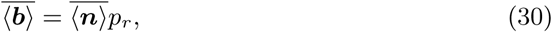

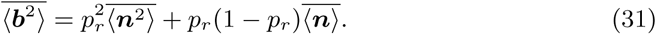

Given formulas for 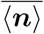 and *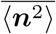* in (18) and (43), respectively, *FF*_*b*_ is computed by applying (30) and (31) in (29). Interestingly, with these formulas one can show the following limits for low (*f* → 0) and high frequency (*f* → ∞) stimulation regardless of ***τ***_*s*_:

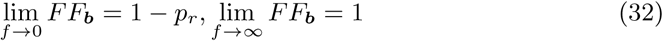

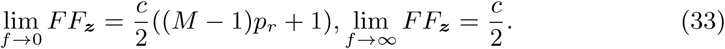

This result assumes that the probability of release *p*_*r*_ is constant. As Fano factor is one for a Poisson distribution, the number of vesicles released follows Poissonian statistics at high frequencies, and is sub-Poissonian at low frequencies. Assuming *c* ≫ 1, the noise in neurotransmitter levels is super-Poissonian at both low and high frequencies.

## 4 Noise Analysis

In this section, we consider specific distributions for the arrival of APs.

### 4.1 Poisson arrival of APs

Let ***τ***_*s*_ be an exponentially-distributed random variable with mean ⟨***τ***_*s*_⟩ = 1*/f*, where *f* is the frequency of AP arrival. Then, as per the moment generating function,

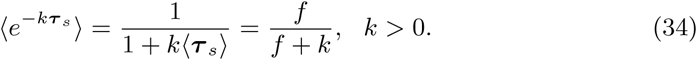

Using (18), (19), (30) and (34), we obtain the following simplified steady-state means

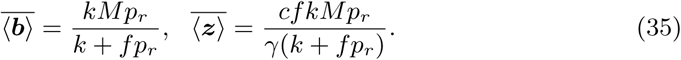

The number of vesicles released per AP decreases with increasing frequency of stimulation due to vesicular depletion in the axon terminal. In contrast, neurotransmitter levels increase monotonically with *f*. Similarly, using (43) and (44), the respective Fano factors can be calculated as

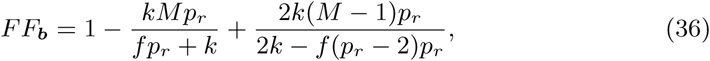

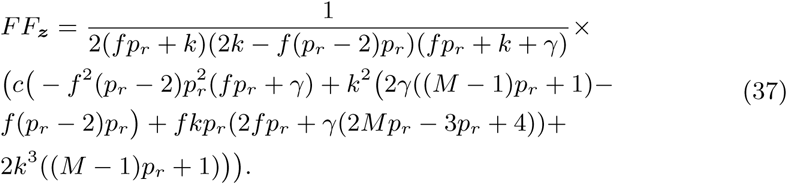

Further analysis reveals an intriguing find: if *Mp*_*r*_ *<* 2, then *FF*_*b*_ monotonically increases with frequency approaching the Poisson limit *FF*_***b***_ *→* 1 as *f → ∞* (Fig. 2). Note in this case *FF*_*b*_ *<* 1 and is always sub-Poissonian. However, if *Mp*_*r*_ *>* 2, then *FF*_*b*_ has a non-monotonic profile, where *FF*_*b*_ *>* 1, i.e., super-Poissonian at an intermediate frequency. Our results further show that *FF*_*z*_ also varies non-monotonically with *f,* but exhibits a minima (Fig. 2).

**Figure 2:**
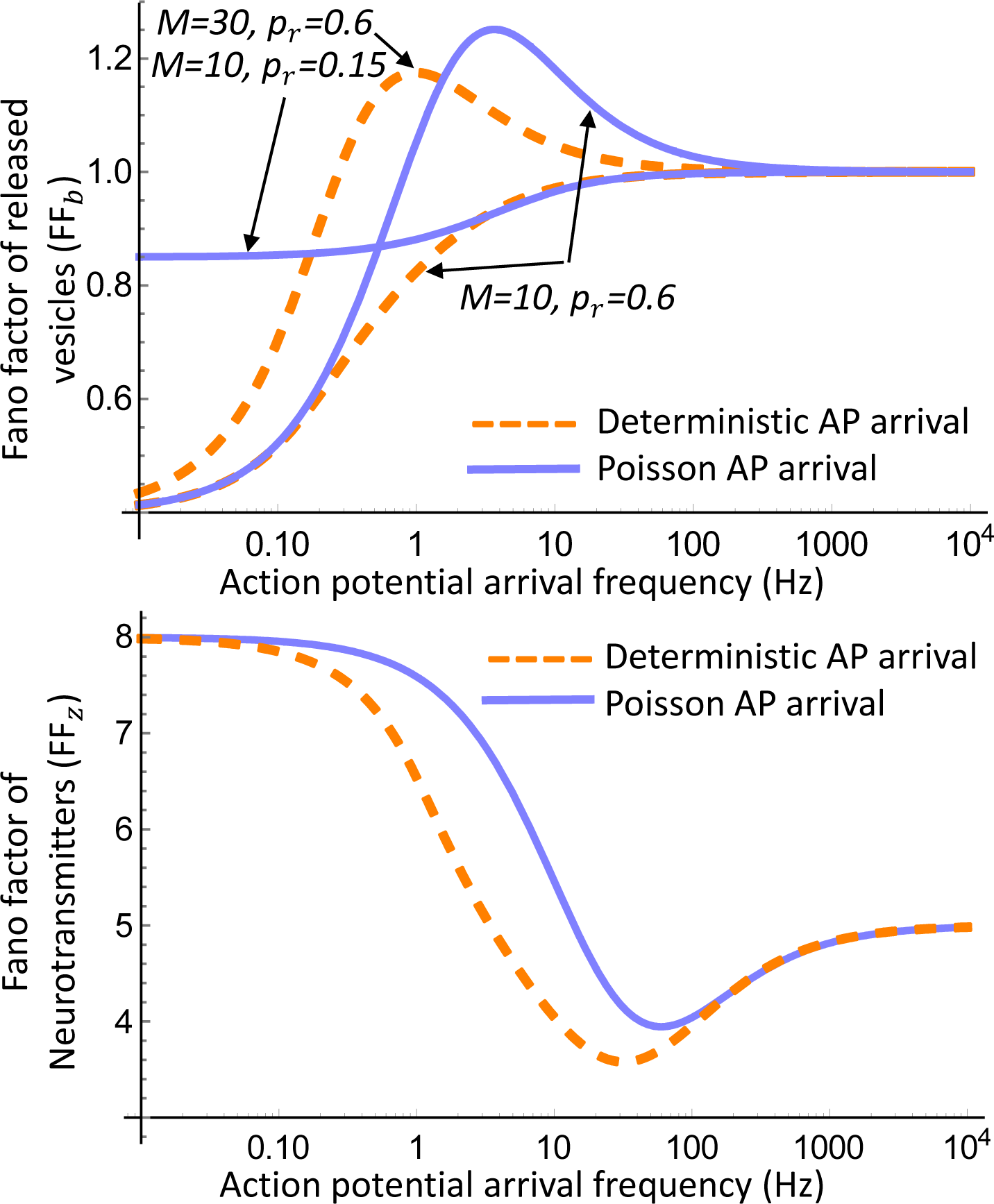
**Top: Noise in the number of released vesicles can be maximized at an intermediate AP arrival frequency.** The Fano factors *FF*_*b*_ in (36) and (38) are plotted as a function of frequency *f* assuming a constant *p*_*r*_ for Poisson and deterministic arrivals. Depending on the values of *M* and *p*_*r*_, the noise can increase monotonically as a function of *f,* or first increase followed by a decrease. See text for exact analytical conditions. For this plot *k* = 1 *sec*^−1^. **Bottom: Noise in the neurotransmitter level is minimized at an intermediate AP arrival frequency**. The Fano factors *FF*_*z*_ in (37) and (39) as a function of frequency *f* showing that while noise is always super-Poissonian (i.e., Fano factor bigger than one), it can be minimal at an optimal frequency. Other parameters are taken as *k* = 3 *sec*^−1^, *γ* = 5 *sec*^−1^, *M* = 5, *c* = 10 and *p*_*r*_ = 0.15.

### 4.2 Deterministic arrival of APs

Next consider a deterministic arrival process where ***τ***_*s*_ = 1*/f* with probability one. In this case, the Fano factors are given by

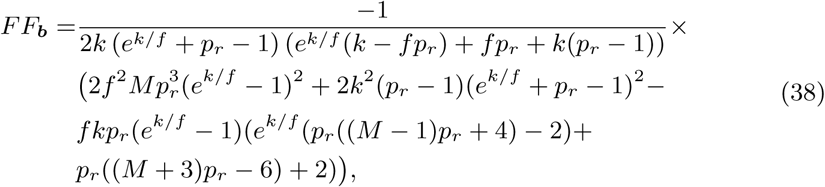

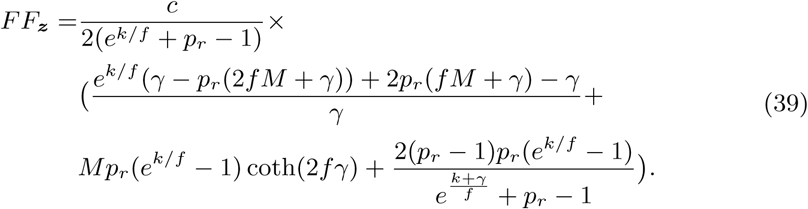

As in the Poisson case, depending on the values of *M* and *p*_*r*_, *FF*_*b*_ can be a monotonic/non-monotonic function of frequency *f* (Fig. 2). More specifically, if 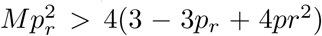, then *FF*_*b*_ is non-monotonic, and monotonic for 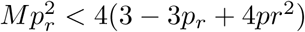.

### 4.3 Frequency-dependent probability of release

Up till now, we have assumed that the probability of release is a constant. We relax this assumption by allowing for a frequency-dependent probability of release. Mechanistically, an increased frequency of APs cause a higher buildup of calcium in the axon terminal which enhances *p*_*r*_. This frequency-dependence is modeled as

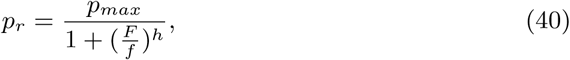

with positive constants *h, p*_*max*_ and *F*. As per this formulation, *p*_*r*_ *→* 0 for *f →* 0, and *p*_*r*_ approaches a maximum value of *p*_*max*_ as *f*. *→ ∞* In the case, the Fano factor limits in (32) and (33) are modified to

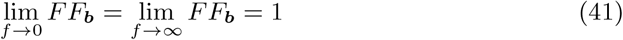

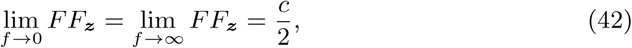

and are the same for low and high-frequency stimulation. While the statistic of vesicle release is Poissonian (*FF*_***b***_ = 1) in the extreme frequency limits, it shows dramatic changes for in-between frequencies with first having a minimum, and then a maximum (Fig. 3). The Fano factor of the neurotransmitter level mirrors these fluctuations with opposite effects – a maximum first followed by a minimum (Fig. 3).

**Figure 3:**
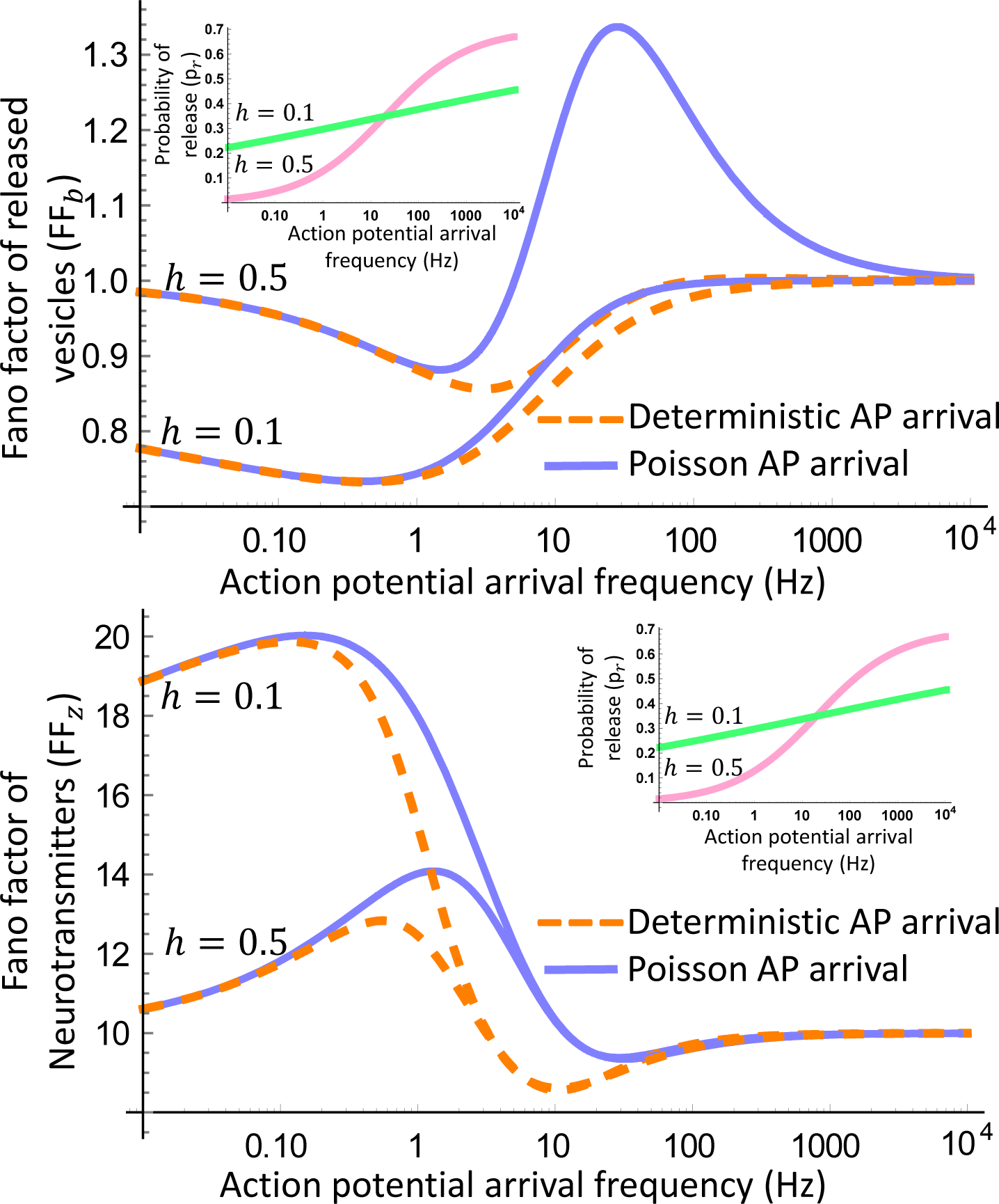
**Top: Noise in the number of released vesicles for a frequency-dependent probability of release**: The inset shows an increase in the probability of release with frequency as per (40). The corresponding *FF*_*b*_ from (36) and (38) are plotted for the same value of *h*, showing noise levels can be both minimal or maximal at intermediate frequencies. In the limit of low and high frequency, *FF*_*b*_ converges to one as per (41). Parameters are taken as *k* = 3 *sec*^−1^, *M* = 30, *F* = 10 *Hz* and *p*_*max*_ = 0.7. **Bottom: Noise in the neurotransmitter level for a frequency-dependent probability of release**: *FF*_*z*_ from (37) and (39) is plotted for frequency-dependent probability of release as shown in the inset. While noise in the neurotransmitter level is always super-Poissonian (assuming *c >>* 1) from (42), it has a non-monotonic profile that can exhibit a minimal or maximal as in the top plot. Parameters are taken as *k* = 1 *sec*^−1^, *γ* = 5 *sec*^−1^, *M* = 5, *c* = 20, *F* = 20 and *p*_*max*_ = 0.7.

## 5 Conclusion

We have applied SHS-based modeling and analysis to systematically characterize the stochastic dynamics of neuronal synaptic transmission. It is important to mention that this contribution also advances the theory for SHS. For example, prior work was focused on either a single timer-dependent reset [33, 46], or two resets where the other occurred at a constant rate [49]. Here we allow the second family of reset to occur at a rate 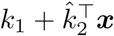 for a constant vector 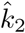 and scalar *k*_1_. Hence, the results on moment derivations presented in Section III are by themselves novel.

Applying the results of Section III to the SHS model in Fig. 1b yields exact analytical formulas for the mean synaptic strength (quantified by (18) and (19)), and the extent of random fluctuations around it (quantified by (43) and (44)). While the noise formulas are quite complicated, they have provided valuable insights in different limits. For example, the limits (32), (33), (41) and (42) show the expected Fano factor at low and high-frequency stimulation for either a constant *p*_*r*_ or frequency-dependent *p*_*r*_. Apart from predicting noise levels, these limits may prove useful for actually inferring parameters by experimentally measuring the corresponding Fano factor, thus providing a unique method that exploits stochastic fluctuations for determining synaptic parameters.

Another interesting finding in the non-monotonicity of noise levels shown in Fig. 2 and Fig.3. For the case of Poisson and deterministic arrivals, we identified parameter regimes that exhibit such behaviors, and these results can be expanded to Gamma or lognormally-distributed AP arrivals. The predictions made in Fig. 2 and Fig.3 can be tested experimentally and, they present an exciting direction of future research in collaboration with neuroscientists.

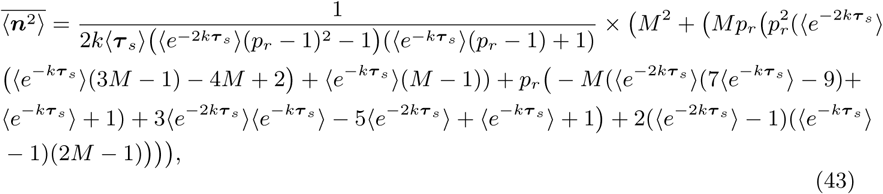

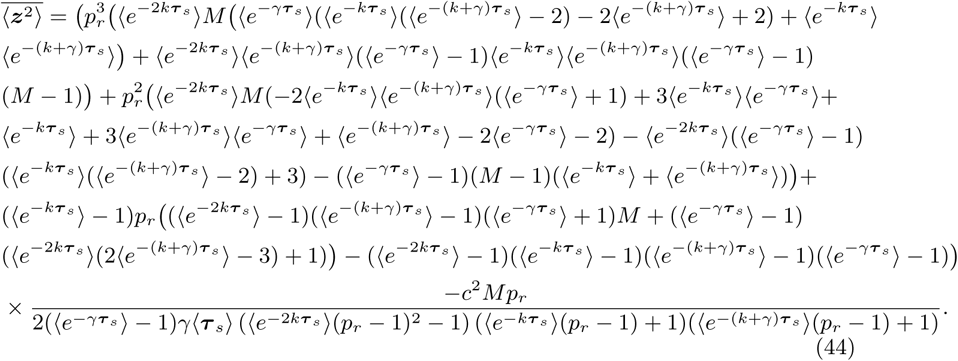

## APPENDIX

In order to prove (14) we use forward Kolmogrov equation as we need to gain the joint probability density function of states ***x*** and timer ***τ***

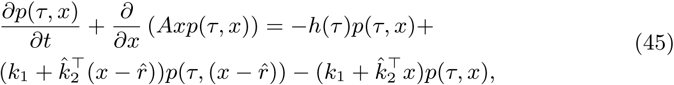

where *∂/∂x* defined as partial derivative vector. Now, having joint probability distribution from (45), conditional mean is

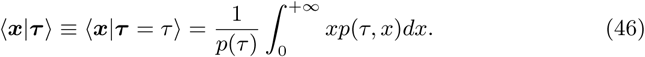

By taking derivative with respect to *t* in (46) we have

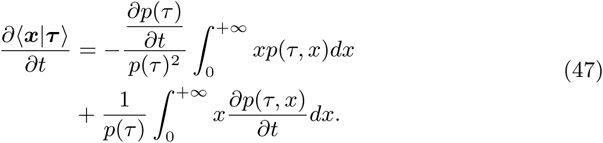

Substituting (5) and (45) into mean dynamics, we have

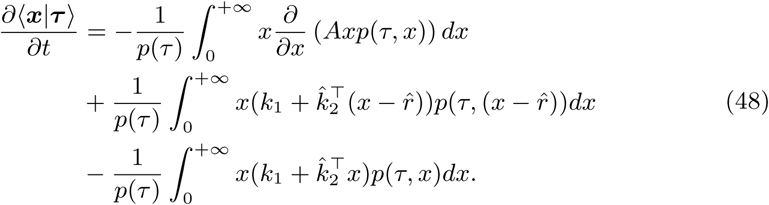

By solving for (48) we get

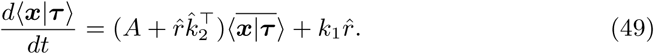

For simplicity, 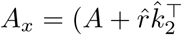 and 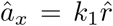 and so we get (14) Similar to (46), conditional second order moment is

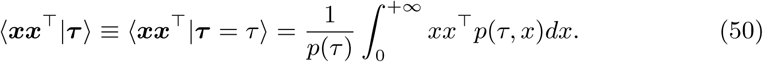

After taking the same steps as (47) and (48) and also also using vectorization [50], 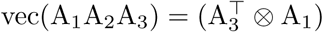 we have

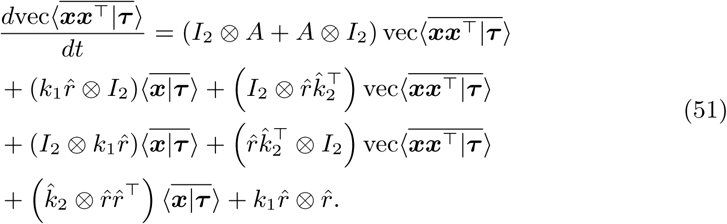

## ACKNOWLEDGMENT

This work is supported by the National Science Foundation Grant ECCS-1711548.

## References

[1] M. V. Tsodyks and H. Markram, “The neural code between neocortical pyramidal neurons depends on neurotransmitter release probability,” Proceedings of the National Academy of Sciences, vol. 94, no. 2, pp. 719–723, 1997.

[2] L. F. Abbott, J. Varela, K. Sen, and S. Nelson, “Synaptic depression and cortical gain control,” Science, vol. 275, no. 5297, pp. 221–224, 1997.

[3] M. Tsodyks, K. Pawelzik, and H. Markram, “Neural networks with dynamic synapses,” Neural computation, vol. 10, no. 4, pp. 821–835, 1998.

[4] W. Senn, H. Markram, and M. Tsodyks, “An algorithm for modifying neurotransmitter release probability based on pre-and postsynaptic spike timing,” Neural Computation, vol. 13, no. 1, pp. 35–67, 2001.

[5] H. Markram, Y. Wang, and M. Tsodyks, “Differential signaling via the same axon of neocortical pyramidal neurons,” Proceedings of the National Academy of Sciences, vol. 95, no. 9, pp. 5323–5328, 1998.

[6] C. Pulido, F. F. Trigo, I. Llano, and A. Marty, “Vesicular release statistics and unitary postsynaptic current at single gabaergic synapses,” Neuron, vol. 85, no. 1, pp. 159–172, 2015.

[7] G. Malagon, T. Miki, I. Llano, E. Neher, and A. Marty, “Counting vesicular release events reveals binomial release statistics at single glutamatergic synapses,” Journal of Neuroscience, vol. 36, no. 14, pp. 4010–4025, 2016.

[8] A. A. Faisal, L. P. Selen, and D. M. Wolpert, “Noise in the nervous system,” Nature reviews neuroscience, vol. 9, no. 4, p. 292, 2008.

[9] F. Chance, “Nelson sb, and abbott lf,” Synaptic depression and the temporal response characteristics of V1 cells. J Neurosci, vol. 18, pp. 4785–4799, 1998.

[10] C. Ribrault, K. Sekimoto, and A. Triller, “From the stochasticity of molecular processes to the variability of synaptic transmission,” Nature Reviews Neuroscience, vol. 12, no. 7, p. 375, 2011.

[11] R. Rosenbaum, J. E. Rubin, and B. Doiron, “Short-term synaptic depression and stochastic vesicle dynamics reduce and shape neuronal correlations,” Journal of neurophysiology, vol. 109, no. 2, pp. 475–484, 2012.

[12] R. Rosenbaum, J. Rubin, and B. Doiron, “Short term synaptic depression with stochastic vesicle dynamics imposes a high-pass filter on presynaptic information,” BMC neuroscience, vol. 13, no. 1, p. O17, 2012.

[13] M. S. Goldman, “Enhancement of information transmission efficiency by synaptic failures,” Neural computation, vol. 16, no. 6, pp. 1137–1162, 2004.

[14] E. Schneidman, B. Freedman, and I. Segev, “Ion channel stochasticity may be critical in determining the reliability and precision of spike timing,” Neural computation, vol. 10, no. 7, pp. 1679–1703, 1998.

[15] C. Zhang and C. S. Peskin, “Improved signaling as a result of randomness in synaptic vesicle release,” Proceedings of the National Academy of Sciences, vol. 112, no. 48, pp. 14 954–14 959, 2015.

[16] A. Arleo, T. Nieus, M. Bezzi, A. D’Errico, E. D’Angelo, and O. J.-M. Coenen, “How synaptic release probability shapes neuronal transmission: information-theoretic analysis in a cerebellar granule cell,” Neural computation, vol. 22, no. 8, pp. 2031–2058, 2010.

[17] C. A. Vargas-García, M. Soltani, and A. Singh, “Conditions for cell size homeostasis: A stochastic hybrid systems approach,” IEEE Life Sciences Letters, vol. 2, pp. 47–50, 2016.

[18] M. Soltani, C. A. Vargas-Garcia, and A. Singh, “Conditional moment closure schemes for studying stochastic dynamics of genetic circuits,” IEEE Transactions on Biomedical Systems and Circuits, vol. 9, pp. 518–526, 2015.

[19] E. Cinquemani, A. Milias-Argeitis, S. Summers, and J. Lygeros, “Stochastic dynamics of genetic networks: modelling and parameter identification,” Bioinformatics, vol. 24, pp. 2748–2754, 2008.

[20] J. Hu, J. Lygeros, and S. Sastry, “Modeling subtilin production in bacillus subtilis using stochastic hybrid systems,” in Hybrid Systems: Computation and Control. Springer, 2004, pp. 417–431.

[21] E. Cinquemani, R. Porreca, G. Ferrari-Trecate, and J. Lygeros, “Subtilin production by bacillus subtilis: Stochastic hybrid models and parameter identification,” IEEE Transactions on Automatic Control, vol. 53, pp. 38–50, 2008.

[22] A. Singh and J. P. Hespanha, “Stochastic hybrid systems for studying biochemical processes,” Philosophical Transactions of the Royal Society A, vol. 368, pp. 4995–5011, 2010.

[23] A. Teel and J. Hespanha, “Stochastic hybrid systems: a modeling and stability theory tutorial,” Proc. of the 54th IEEE Conf. on Decision and Control, Osaka, Japan, 2015.

[24] D. Riley, X. Koutsoukos, and K. Riley, “Modelling and analysis of the sugar cataract development process using stochastic hybrid systems,” IET Systems Biology, vol. 3, pp. 137–154, 2009.

[25] A. Crudu, A. Debussche, and O. Radulescu, “Hybrid stochastic simplifications for multiscale gene networks,” BMC Systems Biology, vol. 3, p. 89, 2009.

[26] D. Antunes and A. Singh, “Computing mRNA and protein statistical moments for a renewal model of stochastic gene-expression,” Proc. of the 52nd IEEE Conf. on Decision and Control, Florence, Italy, pp. 7199–7204, 2013.

[27] J. Pahle, “Biochemical simulations: stochastic, approximate stochastic and hybrid approaches,” Briefings in Bioinformatics, vol. 10, pp. 53–64, 2009.

[28] F. Parise, J. Lygeros, and J. Ruess, “Bayesian inference for stochastic individual-based models of ecological systems: a pest control simulation study,” Environmental Informatics, vol. 3, p. 42, 2015.

[29] K. R. Ghusinga, C. A. Vargas-Garcia, A. Lamperski, and A. Singh, “Exact lower and upper bounds on stationary moments in stochastic biochemical systems,” Physical Biology, vol. 14, p. 04LT01, 2017.

[30] J. Lygeros, K. Koutroumpas, S. Dimopoulos, I. Legouras, P. Kouretas, C. Heichinger, P. Nurse, and Z. Lygerou, “Stochastic hybrid modeling of dna replication across a complete genome,” Proceedings of the National Academy of Sciences, vol. 105, pp. 12 295–12 300, 2008.

[31] C. A. Vargas-García and A. Singh, “Elucidating cell size control mechanisms with stochastic hybrid systems,” IEEE Conference on Decision and Control (CDC), 2018.

[32] K. R. Ghusinga, J. J. Dennehy, and A. Singh, “First-passage time approach to controlling noise in the timing of intracellular events,” Proceedings of the National Academy of Sciences, vol. 114, pp. 693–698, 2017.

[33] M. Soltani and A. Singh, “Moment-based analysis of stochastic hybrid systems with renewal transitions,” Automatica, vol. 84, pp. 62–69, 2017.

[34] E. Buckwar and M. Riedler, “An exact stochastic hybrid model of excitable membranes including spatio-temporal evolution,” Journal of Mathematical Biology, vol. 63, pp. 1051–1093, 2011.

[35] C. Ly and D. Tranchina, “Critical analysis of dimension reduction by a moment closure method in a population density approach to neural network modeling,” Neural Computation, vol. 19, pp. 2032–2092, 2007.

[36] A. Singh, “Modeling noise mechanisms in neuronal synaptic transmission,” bioRxiv, p. 119537, 2017.

[37] A. Singh and J. P. Hespanha, “Stochastic analysis of gene regulatory networks using moment closure,” in Proc. of the 2007 Amer. Control Conference, New York, NY, 2007.

[38] J. P. Hespanha and A. Singh, “Stochastic models for chemically reacting systems using polynomial stochastic hybrid systems,” International Journal of Robust and Nonlinear Control, vol. 15, pp. 669–689, 2005.

[39] A. Singh and J. P. Hespanha, “Approximate moment dynamics for chemically reacting systems,” IEEE Transactions on Automatic Control, vol. 56, pp. 414–418, 2011.

[40] S. M. Ross, “Reliability theory,” in Introduction to Probability Models, 10th ed. Academic Press, 2010, pp. 579–629.

[41] M. Finkelstein, “Failure rate and mean remaining lifetime,” in Failure Rate Modelling for Reliability and Risk, ser. Springer Series in Reliability Engineering. Springer, 2008, pp. 9–44.

[42] M. Soltani and A. Singh, “Stochastic analysis of linear time-invariant systems with renewal transitions,” in American Control Conference, 2017, pp. 1734–1739.

[43] D. Quastel, “The binomial model in fluctuation analysis of quantal neurotransmitter release,” Biophysical Journal, vol. 72, no. 2, pp. 728–753, 1997.

[44] M. Soltani, C. A. Vargas-Garcia, D. Antunes, and A. Singh, “Intercellular variability in protein levels from stochastic expression and noisy cell cycle processes,” PLoS computational biology, vol. 12, no. 8, p. e1004972, 2016.

[45] M. Soltani and A. Singh, “Control design and analysis of a stochastic event-driven system,” IEEE Conference on Decision and Control (CDC), 2018.

[46] D. Antunes, J. P. Hespanha, and C. Silvestre, “Stochastic hybrid systems with renewal transitions: Moment analysis with application to networked control systems with delays,” SIAM Journal on Control and Optimization, vol. 51, pp. 1481–1499, 2013.

[47] M. Soltani and A. Singh, “Moment dynamics for linear time-triggered stochastic hybrid systems,” IEEE 55th Conference on Decision and Control, pp. 3702–3707, 2016.

[48] J. P. Hespanha, “Modeling and analysis of networked control systems using stochastic hybrid systems,” Annual Reviews in Control, vol. 38, pp. 155–170, 2014.

[49] M. Soltani and A. Singh, “Linear piecewise-deterministic markov processes with families of random discrete events,” in European Control Conference, 2018, pp. 447–452.

[50] H. D. Macedo and J. N. Oliveira, “Typing linear algebra: A biproduct-oriented approach,” Science of Computer Programming, vol. 78, no. 11, pp. 2160–2191, 2013.

